# Development of a computational model to inform environmental surveillance sampling plans for *Salmonella enterica* serovar Typhi in wastewater

**DOI:** 10.1101/2023.06.22.545856

**Authors:** Elisabeth Burnor, Cory W. Morin, Jeffry H. Shirai, Nicolette A. Zhou, John Scott Meschke

**Affiliations:** Department of Environmental and Occupational Health Sciences, University of Washington School of Public Health, Seattle, WA, United States of America

**Keywords:** environmental surveillance, pathogen modeling, *Salmonella* Typhi, environmental surveillance (ES), wastewater-based epidemiology (WBE)

## Abstract

Typhoid fever – an acute febrile disease caused by infection with the bacterium *Salmonella enterica* serotype Typhi (*S.* Typhi) – continues to be a leading cause of global morbidity and mortality, especially in developing countries with limited access to safe drinking water and adequate sanitation. Environmental surveillance, the process of detecting and enumerating disease-causing agents in wastewater, enables estimating disease prevalence and trends within a community and is a useful tool to monitor the circulation of typhoid fever in endemic regions. This study presents a computational model, which combines dynamic and probabilistic modeling techniques, to predict – on a spatial and temporal scale – the probability of detecting *S.* Typhi in a wastewater system. This model may be utilized in communities to inform environmental surveillance sampling plans and may provide useful insight into selecting appropriate sampling locations and times.

## 1. Introduction

Typhoid fever is an enteric infectious disease caused by the etiologic agents *Salmonella enterica* serovars Typhi and Paratyphi (*S.* Typhi and *S.* Paratyphi) [1]. Patients with typhoid fever commonly experience prolonged fever, headaches, and abdominal pain [2]. As of 2017, the burden of typhoid fever in low and middle-income countries (LMICs) was estimated to be 17.8 million cases per year, with incidence peaking among children aged 2-4 years old [3]. In 2018 and 2020, the World Health Organization prequalified two typhoid conjugate vaccine (TCV) and has adopted the use of TCVs in parts of the world with a high burden of typhoid [4–6] .Countries considering how and where to provide TCVs need accurate and geographically representative information about typhoid incidence in urban and rural areas [7]. However, the true burden of typhoid is difficult to determine, and data on the incidence of typhoid fever in LMICs remain insufficient [3]. Typhoid fever surveillance is traditionally reliant on clinical-specimen culture surveillance, which requires the detection of *S.* Typhi in blood, bone marrow, stool or urine of individual patients [2]. This type of surveillance is resource-intensive, requires large numbers of participants, and depends on robust laboratory and medical infrastructure [4].

Environmental surveillance (ES) (also termed wastewater-based epidemiology (WBE)) to monitor the circulation of infectious diseases caused by viruses and bacteria has been used to monitor multiple infectious pathogens. These include poliovirus, measles, enteroviruses, hepatitis A, hepatitis E, norovirus, SARS-CoV-2, and others [8]. The COVID-19 pandemic, in particular, brought ES to the attention of the global academic and public health community and SARS-CoV-2 ES and wastewater-based epidemiology has been adopted as a complementary surveillance method to clinical testing for COVID-19 in hundreds of countries [8,9]. ES has also been implemented to monitor the circulation of typhoid fever in endemic geographic areas in order to overcome gaps in clinical population-based surveillance in LMICs [10]. ES offers an anonymous, non-invasive approach to monitor disease circulation in a population, providing the potential for better geographical and temporal resolution for disease prevalence estimates [4,8]. Additionally, a proportion of individuals infected with *S.* Typhi are known to become asymptomatic, chronic carriers, capable of shedding large numbers of bacteria into the environment, but unlikely to receive a clinical test due to lack of symptoms [11,12]. Given this, ES can provide an ideal complementary method to clinical testing for monitoring for potential typhoid transmission and prevalence in a community.

ES strategies for *S.* Typhi present several challenges. It is difficult to identify representative sampling sites in parts of the world that lack centralized wastewater networks [13]. The identification of an optimal sampling time that is both logistically feasible and that captures peak pathogen concentrations and peak probability of detection in wastewater presents an additional challenge [13]. Finally, the sensitivity of a ES sampling plan to detect and quantify pathogens is limited by the sensitivity of the applied laboratory methods [14].

These challenges can be partially addressed by the use of computational modeling. A model was previously designed to quantify the environmental sensitivity of poliovirus ES, given various transmission scenarios, sampling schemes, and laboratory techniques [14]. This model focused on sampling within a single centralized wastewater network and at a single sampling location in Helsinki, Finland, relying on a gamma probability distribution to estimate the delay between fecal output of poliovirus into the wastewater system and its arrival at the sampling site [14]. The authors used this model to estimate the sensitivity of ES with various sampling plans, laboratory method sensitivities, and transmission scenarios. Similarly, a 2020 study presented a different algorithmic approach to augment sampling plans and improve the likelihood of detecting *S.* Typhi in a community with geographic clustering of cases [15]. This study proposed an adaptive sampling site allocation method, where optimal sampling sites are ultimately identified by site performance over time. This study paired the proposed algorithm with an extensive simulation model, used to test the adaptive site allocation method against fixed site locations, in various scenarios [15].

This study takes a different computational approach from these previous works, in order to achieve a similar goal. To inform environmental sampling for the purpose of ES for *S.* Typhi in wastewater, a dynamic computational model is presented with the purpose of predicting the probability of detecting *S.* Typhi at multiple sampling sites within a drainage network and at any hour throughout the day. This model predicts the fate and transport of *S.* Typhi bacteria travelling through a simple but dynamic simulated wastewater system that can be further tailored to assess different kinds of open or closed-channel wastewater networks or drainage systems. Real sewerage networks are likely to be very complex, with branches and structures that may not be well-mapped or documented. This model attempts to simplify these complex systems in order to produce generalized estimates of optimal sampling locations and times. The model then predicts the probability of detecting *S.* Typhi bacteria via environmental sampling and laboratory testing, given pre-specified laboratory method sensitivity.

## 2. Methods

### 2.1 Data Collection

Data for the model were gathered from published literature on bacterial survival dynamics in water and wastewater, estimated wastewater flow per capita, estimated daily fecal output, estimated *S.* Typhi shedding rates in feces, environmental infiltration rates, diurnal variations in wastewater flow, and diurnal variations in fecal output (Source Publication Date Range: 1992 – 2020) (Table S1).

### 2.2 Mapping

In order to run this model, a simple geospatial vector map of the wastewater or drainage network of the community in question is required. Some communities may lack centralized wastewater management systems and associated maps [16,17]. For areas where piped wastewater systems or mapped drainage networks are not available or where wastewater is discharged into environmental streams, a company called Novel-T^©^ has produced an environmental surveillance mapping service (es.world), which provides geospatial maps of synthetic streams and waterways, based on local topology [18]. These maps are designed to identify surveillance sampling sites in streams or drainage systems within a desired catchment area. The simulation provided in this paper utilizes a drainage map shapefile of Vellore City, India, created by Novel-T^©^. The input parameters for Table 1 (see below) were prepared by GIS analysis of this shapefile in R. Briefly, the ‘convergence point’ (the ‘end’ of the system where all drainage lines feed) of the drainage system is identified (see Figure 2). From there, the R script creates a map containing each branch in the system and uses the geospatial coordinates of each branch to determine the parent branches that feed into each branch. This information is incorporated into a spreadsheet file, in which each row defines a single branch, the branch’s parent(s), the length of the branch, the velocity of the wastewater flow, the branch’s input population, and that population’s estimated disease prevalence (see 2.4 Model Function).

### 2.3 Code

The GIS analysis of the drainage map was performed using R v. 4.1.0 [19]. The dynamic computational model was implemented in Python v. 3.7.0 [20].

### 2.4 Model Function

#### 2.4.1 Initialization

Table 1 summarizes the input parameters for the dynamic model that do not change over the course of model simulations. These parameters define each branch of a drainage network and define relationships between branches of the network (*i.e.,* which upstream branches drain into downstream branches). Local information is required to determine the velocity of stream flow (which may be set as constant over the entire system or can be set branch by branch). The user can change fixed parameters for each branch, depending on knowledge about the system and how the population and prevalence of typhoid fever is estimated to be distributed throughout the catchment area. These fixed parameters are then stored in a spreadsheet, which is used during model initialization to create a map of the system. The system map and fixed parameter table may be manually designed by the user. Alternatively, maps may be obtained for multiple cities around the world from es.world (see 2.2 Mapping) [21]. Code is available in the GitHub repository associated with this paper to translate these shapefiles into the fixed parameter inputs in Table 1 (see 2.2 Mapping) or the user may use other GIS software. If no maps or shapefiles exist for a community, the user may manually prepare Table 1 with a visual analysis of open-source satellite imagery or street maps, provided there is some understanding of the direction of flow within a system.

**Table 1:** Fixed Parameter Inputs for Each Branch.

| Input Parameter | Units |
| --- | --- |
| Branch ID | Numerical ID |
| Parent Branches | Numerical IDs for each parent or<br>-1 (for no parent) |
| Velocity | decameters/hour |
| Length (of branch) | decameters |
| Local Population along branch | # persons |
| Estimated Prevalence along branch | Estimated # shedding typhoid cases / total population |

Parameters with defined distributions are shown in Table 2. Values for these parameters are randomly selected from the specified distributions each time a model simulation is performed. The distributions for these parameters were determined by estimates gathered from peer-reviewed literature. All parameters can be adjusted to reflect more accurate, location-specific information.

**Table 2:** Distributed Parameter Inputs for System.

| Input Parameter | Units | Distribution | Values Required |
| --- | --- | --- | --- |
| Fecal Output Rate | grams / day | Log-normal | $\mu, \sigma$ |
| Shedding Rate | pathogen / gram of feces | Simulated distribution | Est. probability of shedding;<br>$\mu, \sigma$ of shedding load |
| Wastewater Flow | Liters / person / day | Log-normal | $\mu, \sigma$ |
| Environmental Flow (including infiltration background flow in a stream) | Liters / hour | Log-normal | $\mu, \sigma$ |
| Bacterial Decay Rate | Proportion remaining after 1 hour | Discrete Uniform | 3 point estimates |

#### 2.4.2 Model Run

Once the system has been initialized, the model will run for *m* days within *n* simulations. Figure 1 presents a flowchart demonstrating the mechanics of the model simulation over days and simulations. Note that each daily update includes data from the previous day in the calculations for the current day (*i.e.,* if a branch of the river has *x* bacteria in it at Hour 24, those bacteria are included in the estimated concentration of bacteria at Hour 0 on the following day). This allows the model to reach ‘steady state’ before calculating the bacterial concentrations in the system for that particular simulation, which is critical, as the model starts from ‘empty’ at the beginning of each simulation. The appropriate number of days, or ‘spin up’ time, to use is a parameter that the user should adjust as needed until the model reliably reaches ‘steady state’ in one simulation. Model plots are available to assess whether this has been achieved. It is important to note that in real transmission scenarios, and in particular, in outbreak scenarios, a ‘steady state’ is unlikely to occur, as the number of new infections and new shedders in the environment and the number of recovered persons who are no longer shedding will change on a daily basis. Although simulating an evolving transmission scenario in conjunction with this model would be an interesting next step, this is beyond the scope of this study.

**Figure 1.**
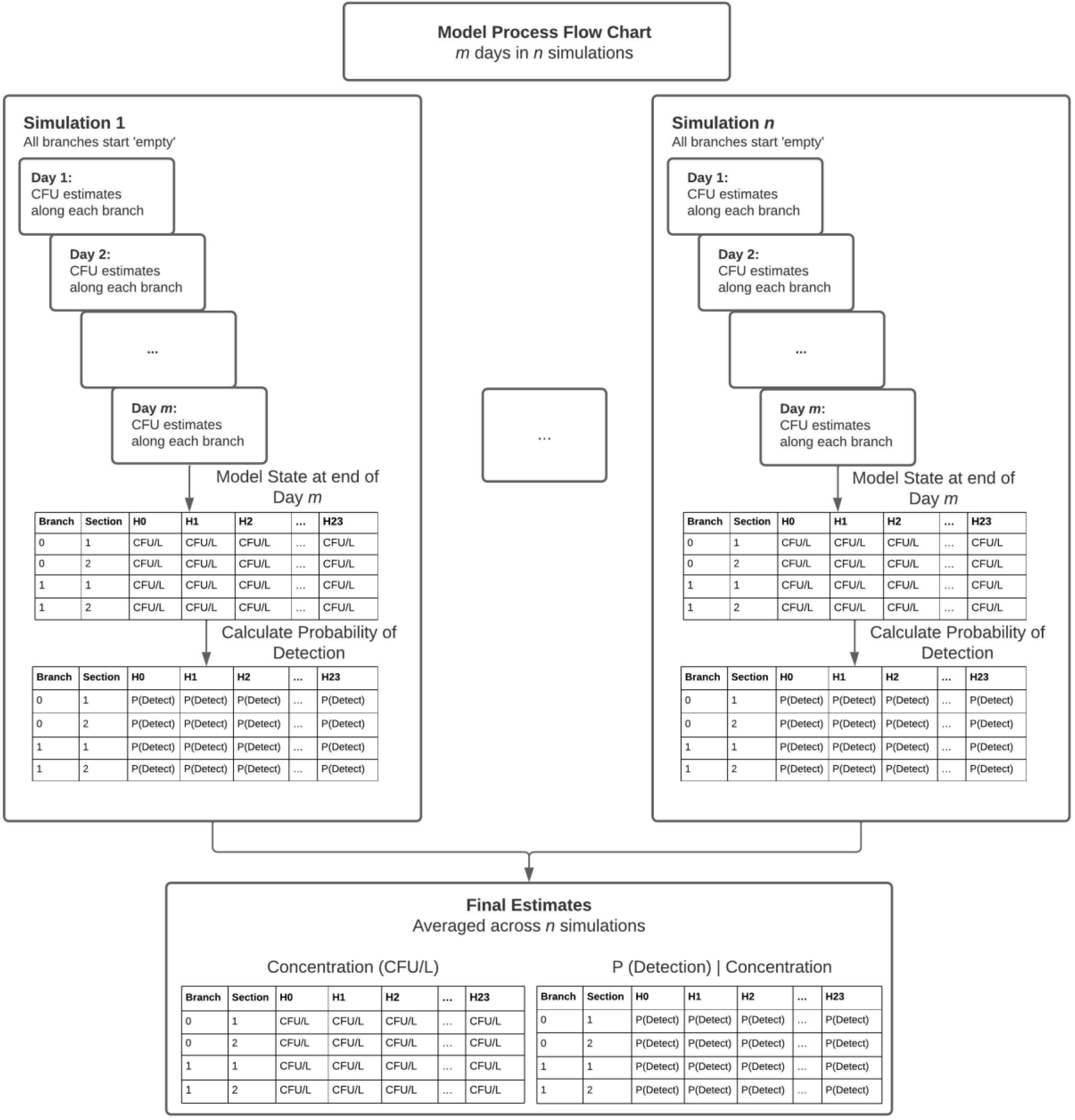
Diagram of the model processing steps. Each simulation box represents one of *n* simulations, for which values for distributed parameters are randomly selected from the distributions pre-specified by the user. Within each simulation, the model iterates over *m* days and stores calculations for bacterial concentrations and probability of detection at the end of day *m*. Once *n* simulations are completed, the model computes averages for bacterial concentrations and probability of detection across all simulations (and associated standard error and confidence intervals).

Wastewater flow and defecation rates are expected to be variable throughout a 24-hour daily cycle. Therefore, the model is designed to update every hour according to the estimated wastewater flow, defecation rates, and environmental infiltration rates at that hour of the day. In the simulation demonstrated below, data were obtained from two studies analyzing variations in wastewater flow over the course of a day and examining hourly variations in defecation rates over a 24-hour cycle [22,23]. These data are available in Tables S2 and S3, but, as wastewater production per capita and defecation rates vary by location and population, it would be preferable to obtain more locally appropriate data for these parameters.

#### 2.4.3 Branch Updates

An example case study is described in the Results section, with the visualization map shown in Figure 2. At every hour, the model starts at the convergence, or ‘end’ of the system (see the ‘Convergence’ point in Figure 2) and calculates pathogen concentration and total flow volume (including wastewater flow volume and infiltration volume) for each section along each branch at that hour. The model follows each branch from the convergence outward by identifying branches and continuing to move from child to parent branch until it reaches a branch that does not have any parent branches (for example, Branches 659 and 891 in Figure 2). Equations and further details on the equations defining pathogen and wastewater flow loading can be found in the supplemental material (See Model Procedure).

**Figure 2.**
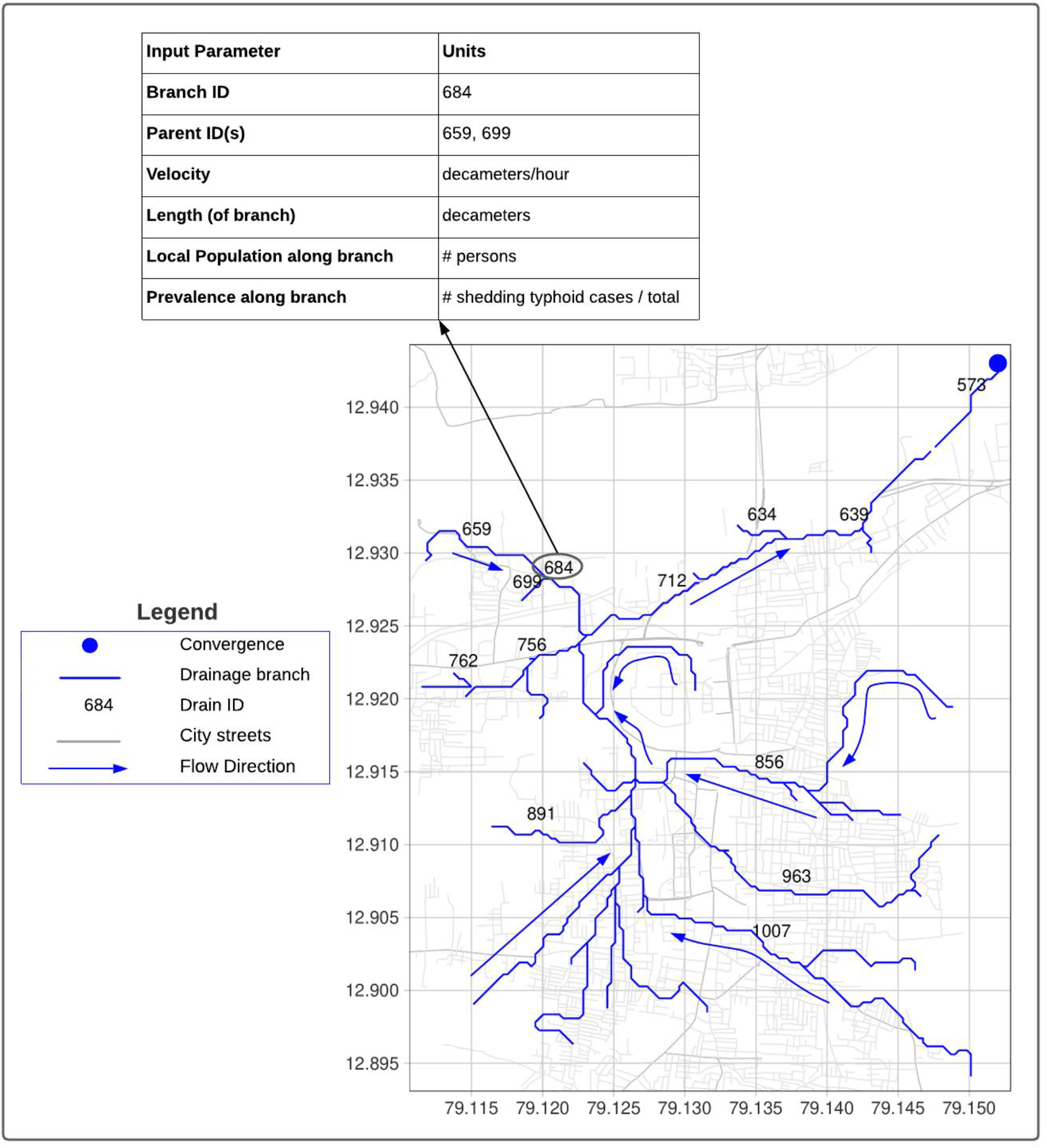
Drainage map of the sewage network of Vellore City, India, identifying the organization and IDs of major drainage branches. This map is used to create the Fixed Parameter table (Table 1), which identifies the length of each branch and each branch’s parent branch(es) in order to represent the direction of wastewater flow. This map provides a visualization of the conceptual map used by the model to track wastewater flow along the sewage network. Blue arrows indicate the direction of flow toward the convergence point. Note: not all branch IDs are shown to improve readability.

#### 2.4.4 Probability of Detection

Once the model has completed *m* days in the n^th^ simulation, it then calculates the probability of detection, given the estimated concentration of bacteria at each section along each branch of the system. The probability of detection is calculated according to the formula described in Ranta, 2001 [14]. These calculations incorporate information about the sensitivity of the laboratory method being used and the volume of wastewater being assayed. Equations and details about these calculations are provided in the supplement.

## 3. Results

To demonstrate an application of this model, a hypothetical scenario was created for Vellore City, India (Figure 2). A drainage map shapefile of the city was obtained from es.world and used to assign ID numbers, parents, length estimates, and population estimates to each branch in order to prepare the parameters described in Table 1 [18]. For this case study, many values for both fixed and distributed variables, such as water velocity, wastewater flow volume, infiltration volume, and fecal shedding rate were estimated using available literature (Supplemental Table S1). These values are not representative of local conditions in Vellore City; thus, the numerical output of the model presented here represents only a simulation of how the model works. In a real-life application, a user would need to incorporate values based on knowledge of the community in question or measurements in order to obtain more accurate model output.

Figure 2 illustrates a visual representation of the case study scenario, where the drainage system of interest is made up of 135 branches, which all feed into ‘Branch 573’. For this scenario, the overall population prevalence of typhoid within this area was set at 200 cases per 100,000 persons and cases were assumed to be uniformly distributed throughout the network. After initialization, the model was run for *m =* 3 days within *n* = 100 simulations.

Figure 3 shows some of the output available after a model run is completed. Figure 3 provides an overview of the entire system over the course of 24 hours, where each bar along the x-axis represents one branch, the bars’ heights on the y-axis correspond to the lengths of the branches, and a heat map shows the probability of detection along each branch, at each hour. Although it is difficult to identify specific segments in this plot, the plot is useful for identifying particular times of day where the probability of detection spikes or dips. To obtain more precise information, the user can specify a branch (according to its ID). Figure 4A shows branch-specific plots, where the x-axis represents hours of the day, the y-axis corresponds to sections along a branch, and a heat map shows the simulated bacterial concentration (left) or the simulated probability of detection (right) at any given section and hour. Figure 4B zooms in further to show the fluctuation in bacterial concentration (left) and probability of detection (right) at a specific section (with a resolution of 10 meters) of a branch, over the course of 24 hours.

**Figure 3.**
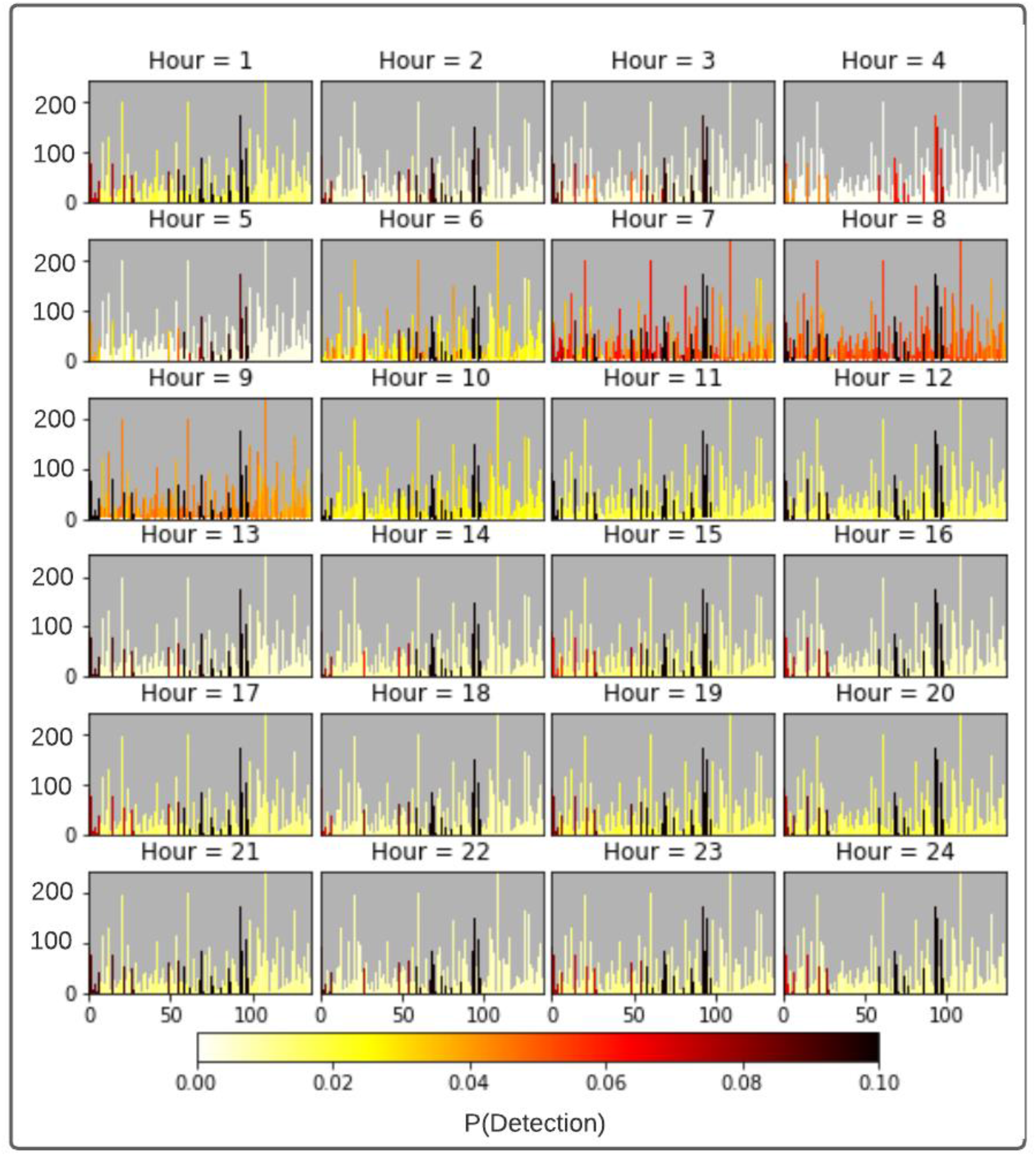
Model output from the case study described in Results. 3A. An hourly plot representing each hour of the day. The x-axis represents each branch in the system (in order of wastewater flow) and the y-axis represents the length of each branch (in meters). A heat map identifies the probability of detection along the length of each branch at each hour of a 24-hour cycle. This plot is helpful in identifying which hours in a 24-hour cycle have the highest probability of detection, across the sewage system.

**Figure 4.**
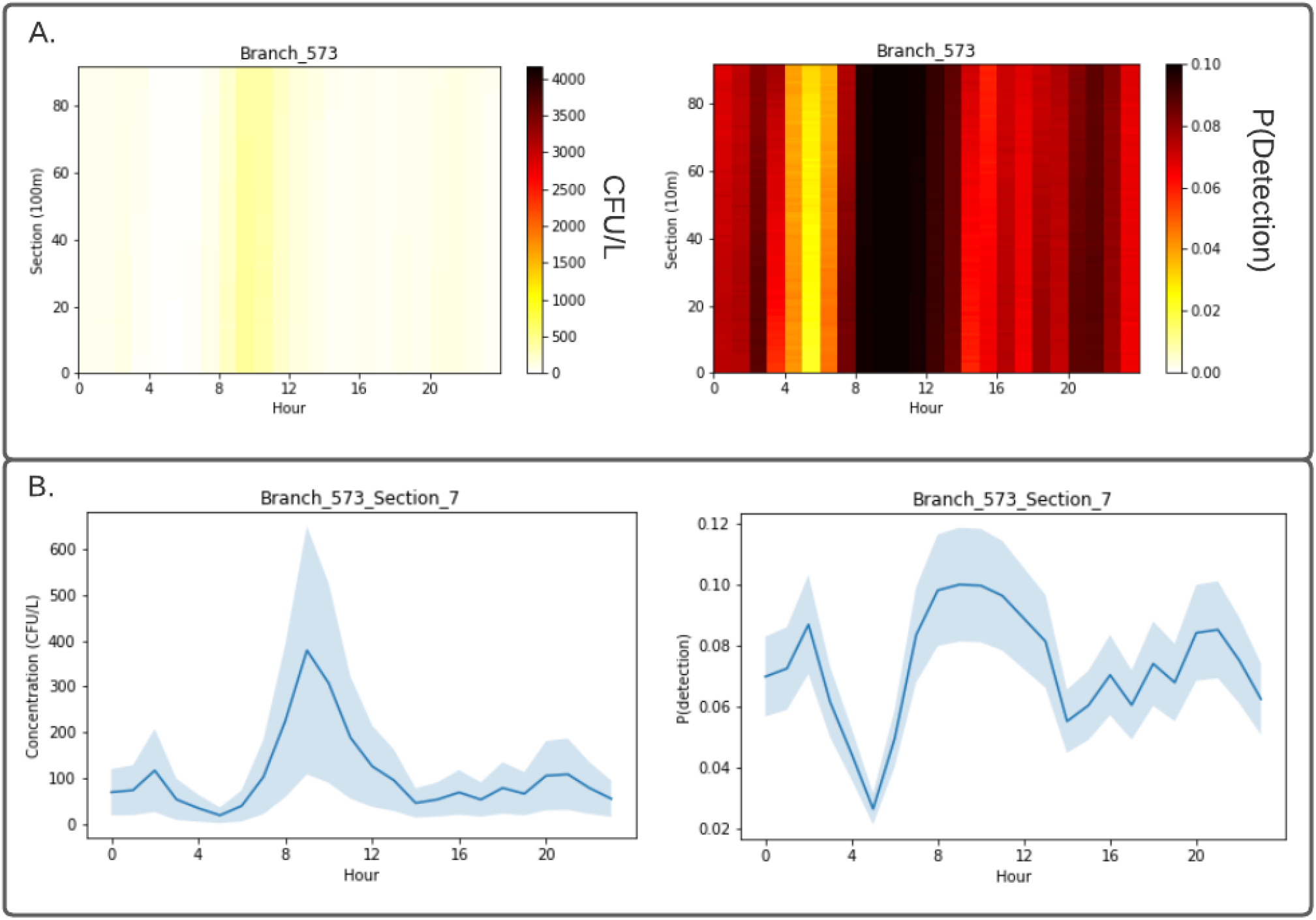
4A. Branch-specific plots for Branch ID 573. Visualizes concentration (left) and probability of detection (right) along each decameter section of a branch at each hour of the day. 4B. Branch- and Section-specific plots for Section 7 of Branch ID 573. Visualizes concentration (left y-axis) and probability of detection (right y-axis) along a single branch, for each hour of the day.

In addition to the plots shown in Figures 3 and 4, the model outputs four spreadsheets (as csv files), estimating the mean concentration of bacteria per liter, standard deviation of these concentrations, mean probability of detection given the bacterial concentration, and standard deviation of these probabilities. All files are in the same format, with raw output data for each branch, section, and hour. Thus, the user can use this raw data to create other plots, assess concentration or probability distributions, or conduct other analyses. See supplement for examples.

## 4. Discussion

The central aim of this model is to track input into and out of sections and branches in a simplified representation of a real-world wastewater drainage system and record how the concentrations and the probability of detection of *S.* Typhi change over time and location. Because of the flexibility of this model, a user has near complete control over all parameters used to describe the system and can adjust these parameters based on local knowledge about wastewater flow rates, drainage line water velocity, and disease prevalence. Because of this flexibility, the accuracy of simulated predictions directly reflect the accuracy of the input data. However, even with limited detailed local information, the model is capable of producing relative estimates for *S.* Typhi concentration and relative probability of detection at sampling locations in or around a city or village within a specified timeframe. Therefore, simulated predictions may provide enough information to help inform decision-making regarding site selection for sampling and the most appropriate time of day to capture samples.

The output plots of the model are useful in a number of ways. Figure 3A is an example of a quick overview of a system, and illustrates, at a glance, whether detection signal from typhoid shedding is present anywhere in the system, or whether it is completely washed out by estimated wastewater volume flow or infiltration. Figures 4A and 4B provide more detailed information that may inform sampling plans. For example, as shown in Figure 4B, there is a clear peak in detection probability between approximately 08:00 and 11:00. However, this timeframe may align with difficult sampling conditions – due to weather, challenging traffic conditions or other factors – and it is useful to know that the probability of detection has smaller peaks at 16:00, 18:00, and 20:00. The branch plots visualized in Figure 4A are created for every branch in the system, and scanning through these plots would allow the user to identify branches with the highest and lowest probabilities of detection, given the local prevalence of typhoid fever. Although this output is from an example simulation, it has some useful implications. Figure 4B illustrates that, although the modelled concentration has a clear morning peak, the probability of detection remains below 20% throughout the day, given the estimated sensitivity of the laboratory method used in this simulation. If an ES program is not detecting *S.* Typhi in locations where the disease is expected to be present, the program might consider continuing to sample in the same location, but increasing detection sensitivity by sampling larger volumes or utilizing a different concentration method [24].

This model also provides considerable flexibility to accommodate different scenarios, drainage systems, local disease prevalence rates, and local disease patterns. The prevalence rates along each branch of the system can be set separately. So, rather than a uniform distribution of disease prevalence across an entire population (which is unlikely to occur), the user could set branches 757 and 782 to have a higher prevalence of typhoid if there is suspicion of a localized cluster of cases in this area. With adjustment of input parameters, such as wastewater flow volume, infiltration, and flow velocity, this model may be applied to any open or closed channel sewer or drainage line network. The model could also be applied to other target pathogens and diseases if the “Organism Decay Rate” and “Shedding Rate” variables in the Distributed Parameters table are adjusted accordingly for different pathogens (Table 2).

It is important to note that the mean wastewater flow velocity used in this example simulation was obtained from estimated wastewater flow velocities in various piped wastewater systems from published literature and do not reflect real-time measured flow velocity estimates, particularly in non-piped systems. The estimated probability of detection of the target pathogen directly relates to the flow and velocity of the stream, open channel, or pipe being sampled. As the model is currently built, the user could increase accuracy by measuring and/or estimating local wastewater flow velocities in portions of the wastewater system. In the future, the overall utility and accuracy of this model could be greatly improved by incorporating accurate regional hydrological data. Models and tools exist in the field of hydrology for estimating hydrological dynamics, and integration of these elements into the existing ES model would be beneficial [25]. This has been done before in microbial risk assessment models assessing complex environmental systems [26]. Portions of the current ES model’s code would need to be adjusted to incorporate this information, but this would be a useful future direction.

There are certain features of the natural and built environment that would have a large impact on a wastewater network, such as pump stations, rainfall, and local industrial waste flows, which are not currently incorporated into this model. It would be useful to add optional pump stations at known locations in order to reflect how those would change the rate of flow in a system and as potential sampling points. The model could also benefit from incorporating variation due to weather and seasonality, as this is likely to change streamflow and background/infiltration flow. For example, Vellore City is situated in the Paler River Basin, which experiences a monsoon season from June to December, during which 80% of the average annual rainfall occurs [27].

Another future consideration for this model would be to test its performance against actual sampling data from an existing ES program for typhoid fever. Currently, such sampling data for *S.* Typhi are unpublished and unavailable. As sampling data from various regions are collected and published, they could be compared against this model’s estimates to evaluate the accuracy of the model’s predictive capability and help identify further areas for improvement within the model code and parameters.

## 5. Conclusion

The model presented here provides the built-in flexibility to utilize both local and published data to provide estimates of pathogen concentrations and detection probabilities at different sampling locations and times. As is the case with any modeling exercise, its output is closely tied to the representativeness and accuracy of the input data. Thus, local information about wastewater flow, streamflow and velocity, and infiltration or background flow will have a large impact on the model’s predictions. Despite challenges in producing accurate absolute estimates, relative estimates provide utility in determining when, where, and how to sample and are also useful in the interpretation of results. Although it is challenging for any model to simulate the complex nature of environmental systems, collaboration with experts in hydrology, meteorology, and environmental science would expand this model’s capacity to better simulate flow dynamics. This in turn would greatly improve the model’s accuracy and flexibility to be adapted to different wastewater systems. Such a tool may be useful for many environmental surveillance scenarios, but it is likely to be particularly useful for communities that lack centralized wastewater networks and treatment systems. Such systems often do not have obvious sampling points, and it may not be clear how sensitive of a laboratory method is required, how many samples to collect, when to collect samples, or which sampling locations will be optimal to capture localized disease clusters. The model presented here is designed to provide a framework to begin answering these questions and a foundation to build upon to produce locally relevant models. The model is relatively simple and very flexible, and has the potential to be adjusted, expanded, and improved to support environmental surveillance efforts by public health practitioners, researchers, and utility operators.

## Supplemental Material

### Model Procedure

At the beginning of each simulation, each branch is ‘empty’, containing zero bacteria and zero liters of water. Given the possibility that some branches may be very long, branches are automatically subdivided into decameter-long sections, and model updates occur for each section of each branch. These updates are performed hourly for both bacterial loading and wastewater flow loading. Supplemental Figure 1 helps visualize this process for Section 1 of one of the branches (Branch 635) of the case study wastewater drainage system described in the main text (Figure 2).

**Supplemental Figure 1:**
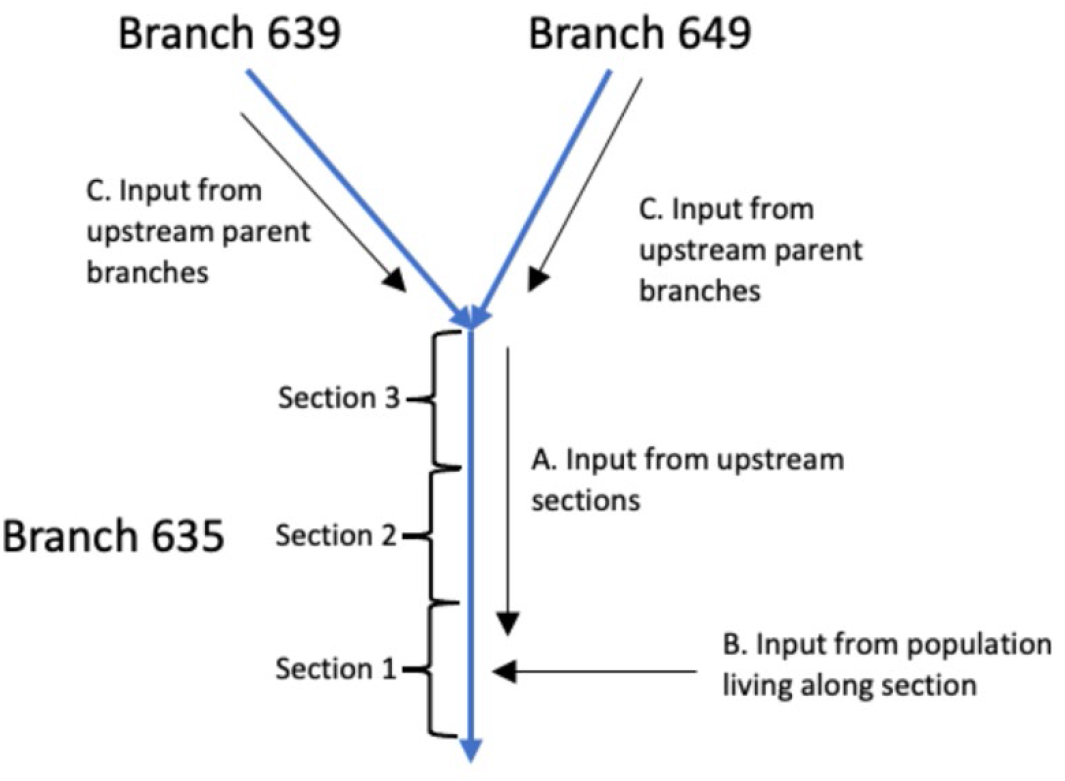
Diagram of the loading process during model estimation of bacterial concentration at Section 1 of Branch 635.

### Model Loading

The number of pathogens (*N*) entering the system at each section of each branch of the system is calculated according to Eq. [1].

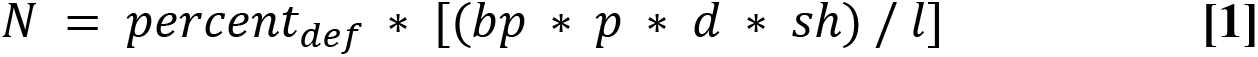

where *percent_def_* is the percent of total daily defecation occurring at the specified hour (Table S2), *bp* is branch population (# of persons), *p* is local prevalence of Typhoid along branch (% of population), *d* is estimated defecation rate (g/day*person), *sh* is shedding rate (CFU / gram), and *l* is length of a branch (meters).

The distance travelled (*D*) by the pathogens and by water flow is calculated according to Eq. [2].

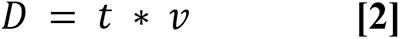

where *t* is time since last update (1 hour) and *v* is velocity of flow in the relevant branch (meters/hour).

The estimated number of pathogens (*P*) in a section of a branch along the system at a given time is calculated by Eq. [3].

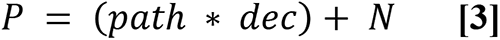

where *path* is the number of pathogens (Colony forming units, or CFUs) feeding into current section from upstream sections (Arrow A in in Figure S1), *dec* is decay rate (hour^-1^), and *N* is the number of pathogens (CFUs) entering the system at that hour from the local population, calculated in Eq. 1 (Arrow B in in Figure S1). If the upstream pathogens feeding into the current section of a branch have travelled farther than the length of that branch (*D,* calculated according to Eq. [2]), the model recursively calculates the cumulative pathogen load from upstream parent branches, given the estimated velocity of each branch and the total distance travelled since the previous hour (Arrows C in in Figure S1).

### Flow Loading

To calculate the total water volume in each section of each branch, the model calculates the input of wastewater flow and background flow (i.e. infiltration in a piped sewage system or streamflow in an open channel or stream system) at each hour of the day. The estimated volume of wastewater and/or environmental water (W) in the system at a given section of a given branch is calculated according to Eq. [4].

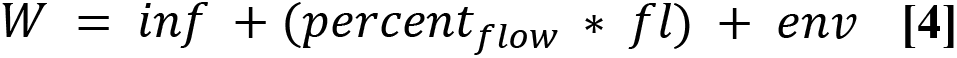

where *inf* (liters) is the influx of wastewater from upstream sections flowing into the current section (Arrow A in in Figure S1), *fl* (liters/hour) is the wastewater flow entering a section of a branch from population living along that branch, *percent_flow_* is the percent of total daily wastewater flow occurring at the specified hour (Table S3), and *env* is the environmental background flow per section of system (liters/hour) (Arrow B in in Figure S1). Similar to the pathogen loading calculations, if the upstream flow entering the current section of a branch has travelled farther than the length of that branch since the previous hour (*D,* calculated according to Eq. [2]), the model recursively calculates the total flow load from all upstream parent branches (Arrows C in Figure S1).

### Pathogen Concentration

Given the pathogen load estimate (P) in Eq [3] and the water volume estimate (W) in Eq [4] at each section of each branch, the model then calculates and stores the estimated concentration (CFU/liter) of *S.* Typhi (C) at each section of each branch at the given hour.

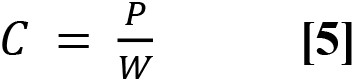

### Probability of Detection

Given laboratory data, practical experience, and/or published literature about the sampling and laboratory detection method of choice, the user provides a value for the probability of a positive result (*P*), given some known number of *S.* Typhi bacteria (*k*) in 1 liter of wastewater. This probability (*P*) is then used to calculate a Beta parameter, which represents the sensitivity of the method; the Beta parameter is used to calculate the probability of a positive result in one-liter samples of wastewater with various estimated concentrations of bacteria.

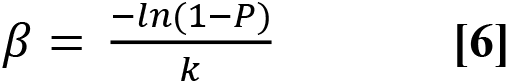

For each section of each branch, the estimated concentration (*C)* of bacteria per liter and a user-specified sampling volume (*S*) (liters) are multiplied to define the shape parameter for a Poisson distribution (*p(b)*) defining the bacterial CFUs in a given sample volume (Eq. [7]).

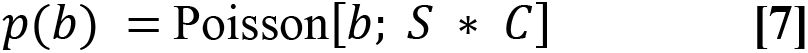

A Monte Carlo simulation with 1,000 simulations is then used to obtain the expected number of *S.* Typhi bacterial CFUs (*b*) (and the mean and standard deviation of the generated probability distribution) captured in a single sample of volume *S,* given repeated random sampling from the Poisson distribution. This expected number of bacteria (*b*) in a collected sample is then used to calculate the probability of a positive laboratory result for *S.* Typhi, via Eq. [8], given the detection method’s Beta parameter (*β*).

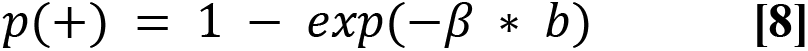

p(+) is the probability of a positive result, *β* is the laboratory method’s sensitivity parameter, as calculated in Eq. [6], and *b* is the expected number of bacteria in the sample.

**Table S1.**
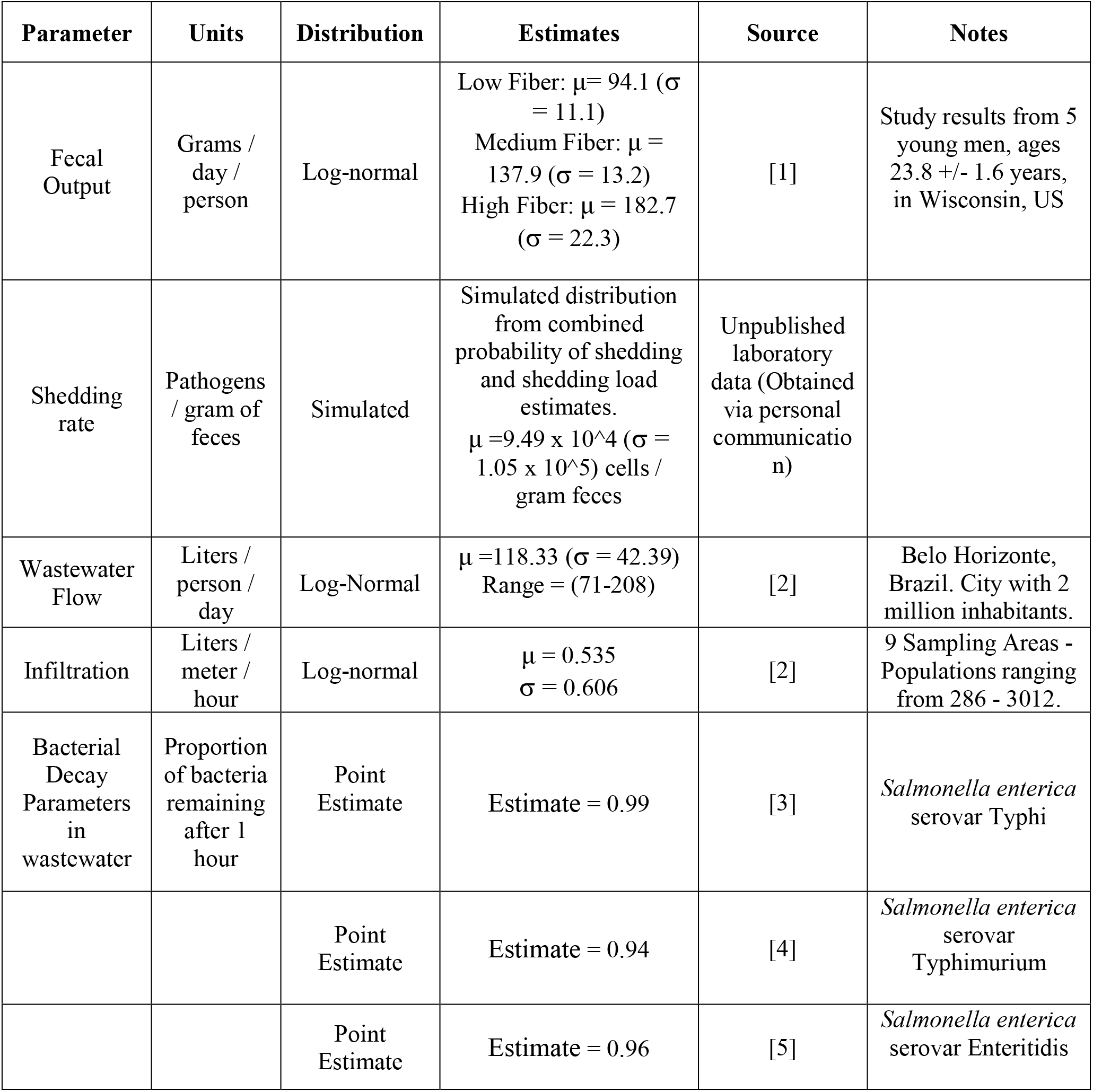
Distributed Parameter Estimates from Literature.

**Table S2:**
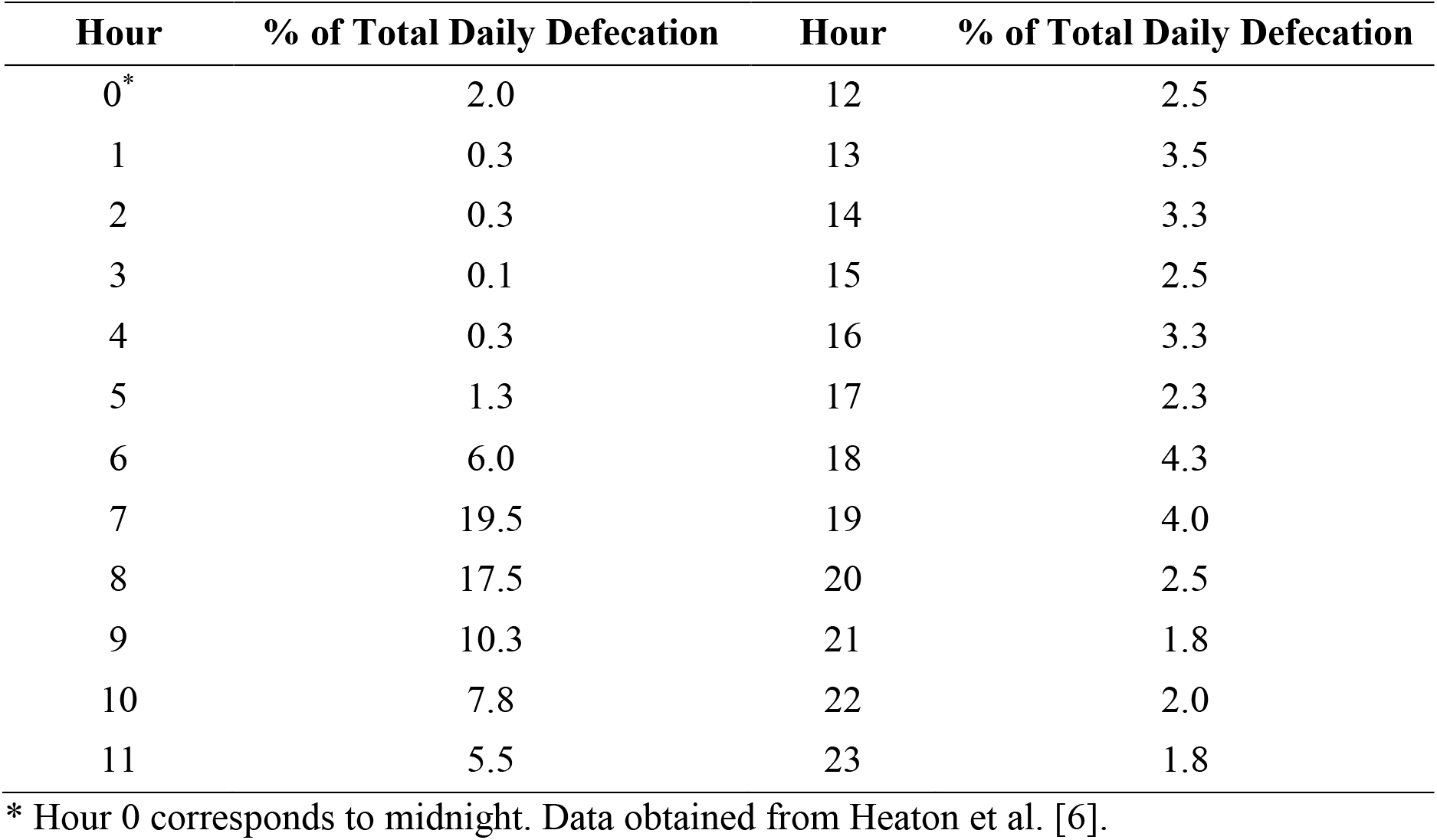
Diurnal Variations in Defecation Rates.

**Table S3:**
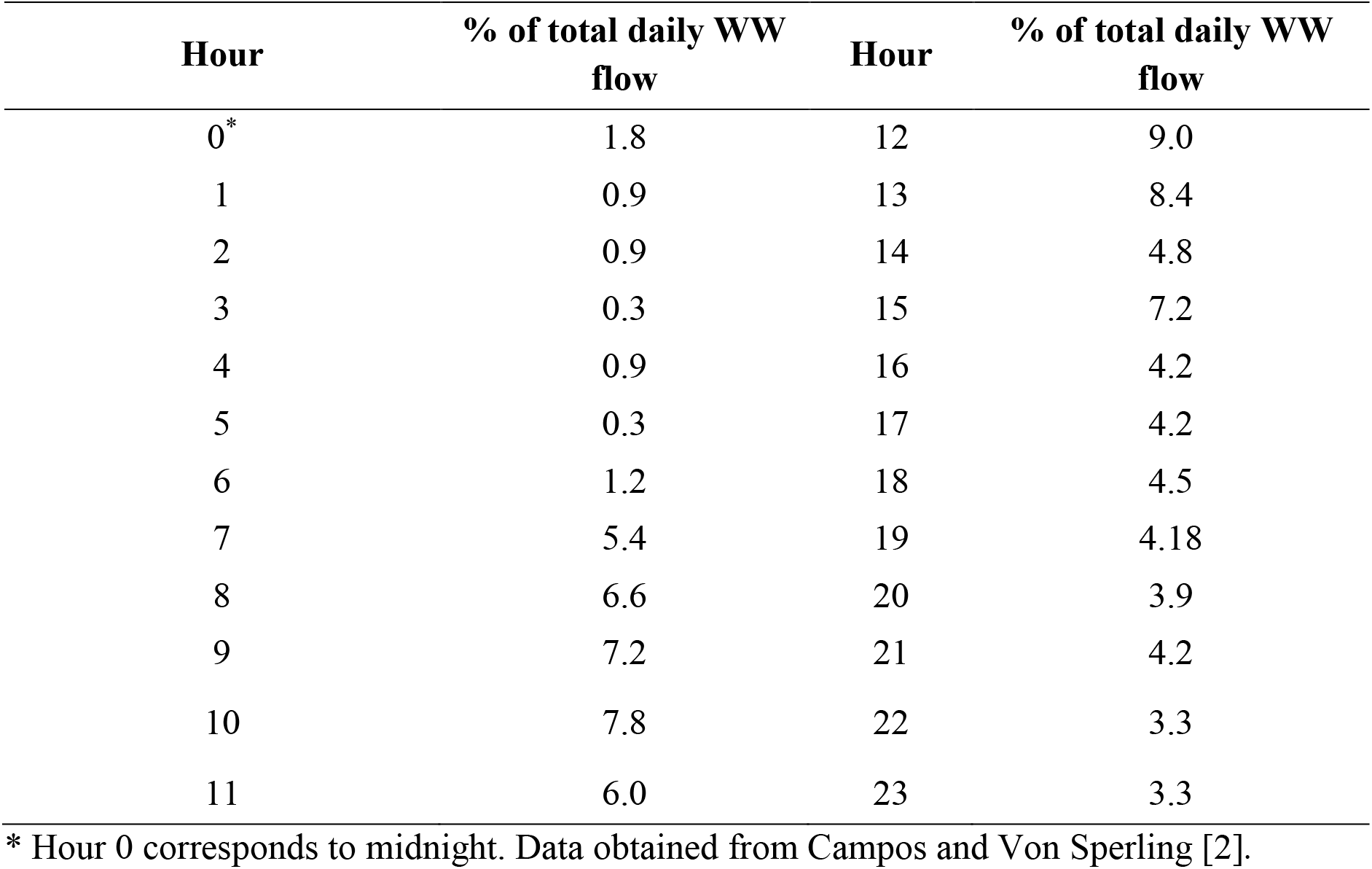
Diurnal Variations in Wastewater (WW) Flow.

## References

1. Akullian A, Ng’eno E, Matheson AI, Cosmas L, Macharia D, Fields B, et al. Environmental Transmission of Typhoid Fever in an Urban Slum. PLoS Negl Trop Dis. 2015 Dec;9(12):e0004212.

2. Su CP, Chen YC, Chang SC. Changing characteristics of typhoid fever in Taiwan. Vol. 37, J Microbiol Immunol Infect. 2004.

3. Antilló M, Warren JL, Crawford FW, Weinberger DM, Kü Rü M E, Pak GD, et al. The burden of typhoid fever in low- and middle-income countries: A meta-regression approach. 2017;

4. Saha S, Tanmoy AM, Andrews JR, Sajib MSI, Yu AT, Baker S, et al. Evaluating PCR-Based Detection of Salmonella Typhi and Paratyphi A in the Environment as an Enteric Fever Surveillance Tool. 2019;100(1):43–6.

5. World Health Organization. Typhoid vaccines: WHO position paper, March 2018 – Recommendations. Vol. 37, Vaccine. Elsevier Ltd; 2019. p. 214–6.

6. World Health Organization. Comparison table of WHO prequalified typhoid conjugate vaccines (TCVs) [Internet]. 2021 [cited 2023 May 31]. Available from: https://apps.who.int/iris/bitstream/handle/10665/345367/WHO-IVB-2021.04-eng.pdf

7. Francy DS, Stelzer EA, Brady AMG, Huitger C, Bushon RN, Ip HS, et al. Comparison of filters for concentrating microbial indicators and pathogens in lake water samples. Appl Environ Microbiol. 2013 Feb;79(4):1342–52.

8. O’keeffe J. Wastewater-based epidemiology: current uses and future opportunities as a public health surveillance tool. 2021;64(3):44–52.

9. Bertels X, Demeyer P, Van den Bogaert S, Boogaerts T, van Nuijs ALN, Delputte P, et al. Factors influencing SARS-CoV-2 RNA concentrations in wastewater up to the sampling stage: A systematic review. Sci Total Environ. 2022 May;820:153290.

10. Saha S, Tanmoy AM, Andrews JR, Sajib MSI, Yu AT, Baker S, et al. Evaluating PCR-Based Detection of Salmonella Typhi and Paratyphi A in the Environment as an Enteric Fever Surveillance Tool. Am J Trop Med Hyg. 2019;100(1):43.

11. Ames WR, Robins M. Age and Sex as Factors in the Development of the Typhoid Carrier State, and a Method for Estimating Carrier Prevalence. Am J Public Health Nations Health. 2008;33(3):221–30.

12. Gopinath S, Carden S, Monack D. Shedding light on Salmonella carriers. Trends Microbiol. 2012;20(7):320–7.

13. Hovi T, Shulman LM, van der Avoort H, Deshpande J, Roivainen M, De Gourville EM. Role of environmental poliovirus surveillance in global polio eradication and beyond. Epidemiol Infect. 2012 Jan;140(1):1–13.

14. Ranta J, Hovi T, Arjas E. Poliovirus surveillance by examining sewage water specimens: Studies on detection probability using simulation models. Risk Anal. 2001;21(6):1087–96.

15. Wang Y, Moe CL, Dutta S, Wadhwa A, Kanungo S, Mairinger W, et al. Designing a typhoid environmental surveillance study: A simulation model for optimum sampling site allocation. Epidemics. 2020 Jun 1;31:100391.

16. Ali S, Gudina EK, Gize A, Aliy A, Adankie BT, Tsegaye W, et al. Community Wastewater-Based Surveillance Can Be a Cost-Effective Approach to Track COVID-19 Outbreak in Low-Resource Settings: Feasibility Assessment for Ethiopia Context. Int J Environ Res Public Health. 2022 Jul 1;19(14).

17. Uzzell CB, Troman CM, Rigby J, Raghava Mohan V, John J, Abraham D, et al. Environmental surveillance for Salmonella Typhi as a tool to estimate the incidence of typhoid fever in low-income populations. Wellcome Open Res. 2023 Jan 6;8:9.

18. Novel-T. Environmental Surveillance: Supporting Polio Eradication [Internet]. es.world. [cited 3030 May 30]. Available from: https://es.world/

19. R Core Team. R: A language and environment for statistical computing. [Internet]. Vienna, Austria: R Foundation for Statistical Computing; 2021. Available from: https://www.R-project.org/

20. Van Rossum, G, Drake, F. L. Python 3 Reference Manual. CreateSpace.; 2009.

21. Novel-T. Environmental Surveillance: Supporting Polio Eradication [Internet]. [cited 2020 May 3]. Available from: es.world

22. Heaton KW, Radvan J, Cripps H, Mountford RA, Braddon FE, Hughes AO. Defecation frequency and timing, and stool form in the general population: a prospective study. Gut. 1992 Jun;33(6):818–24.

23. Campos HM, von Sperling M. Estimation of domestic wastewater characteristics in a developing country based on socio-economic variables. Water Sci Technol. 1996 Jan 1;34(3):71–7.

24. Zhou N, Ong A, Fagnant-Sperati C, Harrison J, Kossik A, Beck N, et al. Evaluation of Sampling and Concentration Methods for Salmonella enterica Serovar Typhi Detection from Wastewater. Am J Trop Med Hyg. 2023 Mar 1;108(3):482–91.

25. Eregno FE, Tryland I, Tjomsland T, Myrmel M, Robertson L, Heistad A. Quantitative microbial risk assessment combined with hydrodynamic modelling to estimate the public health risk associated with bathing after rainfall events. Sci Total Environ. 2016 Apr 1;548– 549:270–9.

26. Clark CS. Potential and Actual Biological Related Health Risks of Wastewater Industry Employmen. Water Pollut Control Fed. 1987;59(12):999–1008.

27. Fan X, Luo X. Precipitation and flow variations in the Lancang-Mekong River Basin and the implications of monsoon fluctuation and regional topography. Water Switz. 2019;11(10).

## References

1. Kurasawa S, Haack VS, Marlett JA. Plant residue and bacteria as bases for increased stool weight accompanying consumption of higher dietary fiber diets. J Am Coll Nutr. 2000 Aug;19(4):426–33.

2. Campos HM, Von Sperling M. Estimation of domestic wastewater characteristics in a developing country based on socio-economic variables. Vol. 34, Water Science and Technology. 1996. p. 71–7.

3. Cho JC, Kim SJ. Viable, but non-culturable, state of a green fluorescence protein-tagged environmental isolate of Salmonella typhi in groundwater and pond water. FEMS Microbiol Lett. 1999 Jan 1;170(1):257–64.

4. Himathongkham S, Bahari S, Riemann H, Cliver D. Survival of Escherichia coli O157:H7 and Salmonella typhimurium in cow manure and cow manure slurry. FEMS Microbiol Lett. 1999 Sep 15;178(2):251–7.

5. Ravva SV, Sarreal CZ. Survival of Salmonella enterica in aerated and nonaerated wastewaters from dairy lagoons. Int J Environ Res Public Health. 2014 Oct 29;11(11):11249–60.

6. Heaton KW, Radvan J, Cripps H, Mountford RA, Braddon FEM, Hughes A 0. Defecation frequency and timing, and stool form in the general population: a prospective study. Gut. 1992;33:818–24.

