## supplemental material for "Development of a computational model to inform environmental surveillance sampling plans for *Salmonella enterica* serovar Typhi in wastewater"

Short title: Typhoid environmental surveillance model

Elisabeth Burnor[1], Cory W. Morin[1], Jeffry H. Shirai[1], Nicolette A. Zhou[1], John Scott Meschke[1*]

**Affiliation:**

[1] Department of Environmental and Occupational Health Sciences, University of Washington School of Public Health, Seattle, WA, United States of America

* Corresponding author
 (JSM)

### Model Procedure

At the beginning of each simulation, each branch is 'empty', containing zero bacteria and zero liters of water. Given the possibility that some branches may be very long, branches are automatically subdivided into decameter-long sections, and model updates occur for each section of each branch. These updates are performed hourly for both bacterial loading and wastewater flow loading. Supplemental Figure 1 helps visualize this process for Section 1 of one of the branches (Branch 635) of the case study wastewater drainage system described in the main text (Figure 2).

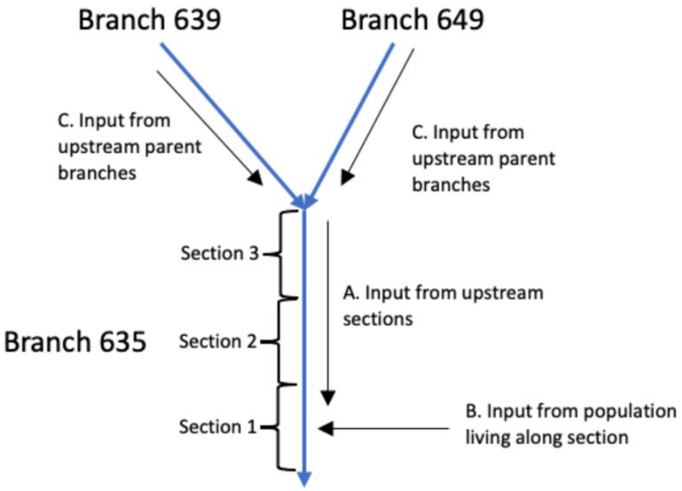

**Supplemental Figure 1:** Diagram of the loading process during model estimation of bacterial concentration at Section 1 of Branch 635.

### Model Loading

The number of pathogens ($N$) entering the system at each section of each branch of the system is calculated according to Eq. [1].

$$N = percent_{def} * [(bp * p * d * sh) / l] \quad \textbf{[1]}$$

where $percent_{def}$ is the percent of total daily defecation occurring at the specified hour (Table S2), *bp* is branch population (# of persons), *p* is local prevalence of Typhoid along branch (% of population), *d* is estimated defecation rate (g/day*person), *sh* is shedding rate (CFU / gram), and *l* is length of a branch (meters).

The distance travelled (*D*) by the pathogens and by water flow is calculated according to Eq. [2].

$$D = t * v \qquad \textbf{[2]}$$

where *t* is time since last update (1 hour) and *v* is velocity of flow in the relevant branch (meters/hour).

$$p(+) = 1 - exp(-\beta * b) \qquad \textbf{[8]}$$

p(+) is the probability of a positive result, $\beta$ is the laboratory method's sensitivity parameter, as calculated in Eq. [6], and *b* is the expected number of bacteria in the sample.

**Table S1. Distributed Parameter Estimates from Literature**

| Parameter | Units | Distribution | Estimates | Source | Notes |
|---|---|---|---|---|---|
| Fecal Output | Grams / day / person | Log-normal | Low Fiber: $\mu$= 94.1 ($\sigma$ = 11.1) Medium Fiber: $\mu$ = 137.9 ($\sigma$ = 13.2) High Fiber: $\mu$ = 182.7 ($\sigma$ = 22.3) | [1] | Study results from 5 young men, ages 23.8 +/- 1.6 years, in Wisconsin, US |
| Shedding rate | Pathogens / gram of feces | Simulated | Simulated distribution from combined probability of shedding and shedding load estimates. $\mu$ =9.49 x 10^4 ($\sigma$ = 1.05 x 10^5) cells / gram feces | Unpublished laboratory data (Obtained via personal communicatio n) | |
| Wastewater Flow | Liters / person / day | Log-Normal | $\mu$ =118.33 ($\sigma$ = 42.39) Range = (71-208) | [2] | Belo Horizonte, Brazil. City with 2 million inhabitants. |
| Infiltration | Liters / meter / hour | Log-normal | $\mu$ = 0.535 $\sigma$ = 0.606 | [2] | 9 Sampling Areas - Populations ranging from 286 - 3012. |
| Bacterial Decay Parameters in wastewater | Proportion of bacteria remaining after 1 hour | Point Estimate | Estimate = 0.99 | [3] | *Salmonella enterica* serovar Typhi |
| | | Point Estimate | Estimate = 0.94 | [4] | *Salmonella enterica* serovar Typhimurium |
| | | Point Estimate | Estimate = 0.96 | [5] | *Salmonella enterica* serovar Enteritidis |

**Table S2:** Diurnal Variations in Defecation Rates

| Hour | % of Total Daily Defecation | Hour | % of Total Daily Defecation |
|---|---|---|---|
| 0* | 2.0 | 12 | 2.5 |
| 1 | 0.3 | 13 | 3.5 |
| 2 | 0.3 | 14 | 3.3 |
| 3 | 0.1 | 15 | 2.5 |
| 4 | 0.3 | 16 | 3.3 |
| 5 | 1.3 | 17 | 2.3 |
| 6 | 6.0 | 18 | 4.3 |
| 7 | 19.5 | 19 | 4.0 |
| 8 | 17.5 | 20 | 2.5 |
| 9 | 10.3 | 21 | 1.8 |
| 10 | 7.8 | 22 | 2.0 |
| 11 | 5.5 | 23 | 1.8 |

* Hour 0 corresponds to midnight. Data obtained from Heaton et al. [6].

**Table S3:** Diurnal Variations in Wastewater (WW) Flow

| Hour | % of total daily WW flow | Hour | % of total daily WW flow |
|---|---|---|---|
| 0* | 1.8 | 12 | 9.0 |
| 1 | 0.9 | 13 | 8.4 |
| 2 | 0.9 | 14 | 4.8 |
| 3 | 0.3 | 15 | 7.2 |
| 4 | 0.9 | 16 | 4.2 |
| 5 | 0.3 | 17 | 4.2 |
| 6 | 1.2 | 18 | 4.5 |
| 7 | 5.4 | 19 | 4.18 |
| 8 | 6.6 | 20 | 3.9 |
| 9 | 7.2 | 21 | 4.2 |
| 10 | 7.8 | 22 | 3.3 |
| 11 | 6.0 | 23 | 3.3 |

* Hour 0 corresponds to midnight. Data obtained from Campos and Von Sperling [2].

### References

1. Kurasawa S, Haack VS, Marlett JA. Plant residue and bacteria as bases for increased stool weight accompanying consumption of higher dietary fiber diets. J Am Coll Nutr. 2000 Aug;19(4):426–33.

2. Campos HM, Von Sperling M. Estimation of domestic wastewater characteristics in a developing country based on socio-economic variables. Vol. 34, Water Science and Technology. 1996. p. 71–7.

3. Cho JC, Kim SJ. Viable, but non-culturable, state of a green fluorescence protein-tagged environmental isolate of Salmonella typhi in groundwater and pond water. FEMS Microbiol Lett. 1999 Jan 1;170(1):257–64.

4. Himathongkham S, Bahari S, Riemann H, Cliver D. Survival of Escherichia coli O157:H7 and Salmonella typhimurium in cow manure and cow manure slurry. FEMS Microbiol Lett. 1999 Sep 15;178(2):251–7.

5. Ravva SV, Sarreal CZ. Survival of Salmonella enterica in aerated and nonaerated wastewaters from dairy lagoons. Int J Environ Res Public Health. 2014 Oct 29;11(11):11249–60.

6. Heaton KW, Radvan J, Cripps H, Mountford RA, Braddon FEM, Hughes A 0. Defecation frequency and timing, and stool form in the general population: a prospective study. Gut. 1992;33:818–24.
